# A Human Liver Organoid Screening Platform for DILI Risk Prediction

**DOI:** 10.1101/2021.08.26.457824

**Authors:** Charles J. Zhang, Sophia R. Meyer, Matthew J. O’Meara, Sha Huang, Meghan M. Capeling, Daysha Ferrer-Torres, Charlie J. Childs, Jason R. Spence, Robert J. Fontana, Jonathan Z. Sexton

**Affiliations:** Department of Medicinal Chemistry, College of Pharmacy, University of Michigan, Ann Arbor, MI, 48109, USA; Department of Computational Medicine and Bioinformatics, University of Michigan, Ann Arbor, MI, 48109, USA; Department of Internal Medicine, Gastroenterology and Hepatology, Michigan Medicine at the University of Michigan, Ann Arbor, MI, 48109, USA; Department of Cell and Developmental Biology, University of Michigan, Ann Arbor, MI, 48109, USA; U-M Center for Drug Repurposing, University of Michigan, Ann Arbor, MI, 48109, USA

**Keywords:** microfluidic devices, drug development, hepatotoxicity, high-content imaging

## Abstract

**Background and Aims:** Drug-induced liver injury (DILI), both intrinsic and idiosyncratic, causes frequent morbidity, mortality, clinical trial failures and post-approval withdrawal. This suggests an unmet need for improved in vitro models for DILI risk prediction that can account for diverse host genetics and other clinical factors. In this study, we evaluated the utility of human liver organoids (HLOs) for high-throughput DILI risk prediction and in an organ-on-chip system.

**Methods:** HLOs were derived from 3 separate iPSC lines and benchmarked on two platforms for their ability to model *in vitro* liver function and identify hepatotoxic compounds using biochemical assays for albumin, ALT, and AST, microscopy-based morphological profiling, and single-cell transcriptomics: 1) HLOs dispersed in 384-well formatted plates and exposed to a library of compounds. 2) HLOs adapted to a liver-on-chip system.

**Results:**

1. Dispersed HLOs derived from the 3 iPSC lines had similar DILI predictive capacity to intact HLOs in a high-throughput screening format allowing for measurable IC50 values of compound cytotoxicity. Distinct morphological differences were observed in cells treated with drugs exerting differing mechanisms of toxicity.
2. On-chip HLOs significantly increased albumin production, CYP450 expression, and ALT/AST release when treated with known DILI drugs compared to dispersed HLOs and primary human hepatocytes. On-chip HLOs were able to predict the synergistic hepatotoxicity of tenofovir-inarigivir and showed steatosis and mitochondrial perturbation via phenotypic and transcriptomic analysis with exposure to FIAU and acetaminophen, respectively.

**Conclusions:** The high throughput and liver-on-chip system exhibit enhanced *in vivo*-like function and demonstrate the potential utility of these platforms for hepatotoxicity risk assessment. Tenofovir-inarigivr associated hepatotoxicity was observed and correlates with the clinical manifestation of DILI observed in patients.

**LAY SUMMARY:** Idiosyncratic (spontaneous, patient-specific) drug-induced liver injury (DILI) is difficult to study due to the lack of liver models that function as human liver tissue and are adaptable for large-scale drug screening. Human liver organoids grown from patient stem cells respond to known DILI-causing drugs in both a high-throughput and on a physiological “chip” culture system. These platforms show promise in their use as predictive model for novel drugs before entering clinical trials.

## INTRODUCTION

Drug-induced liver injury (DILI) is an infrequent but important cause of both acute and chronic liver disease.[1,2] An estimated 22% of clinical trial failures and 32% of market withdrawals of therapeutics are due to hepatotoxicity.[3,4] Hepatotoxicity is typically not identified until clinical trials or post-marketing which creates an increased risk for clinical trial participants as well as a financial burden in drug development. Most instances of DILI are termed “idiosyncratic,” since they are largely independent of the dose and duration of drug use and develop in only a small proportion of treated patients for as-yet unclear reasons. As there are currently over 1000 prescription medications and >80,000 herbal and dietary supplements (HDS) available for use in the United States [5], the potential for additive or synergistic liver toxicity is high with low predictive capability.

Recently, inarigivir soproxil (GS-9992) was investigated against Hepatitis B (HBV).[6] Inarigivir monotherapy and in combination with tenofovir showed no clear signs of toxicity in two clinical trials for HBV. However, a later phase-2 inarigivir/tenofovir study identified severe DILI in patients given the combination of both drugs after 16 weeks of therapy that lead to the discontinuation of this drug development program. All 7 patients had an elevated alanine aminotransferease (ALT) after 16 weeks of therapy, 4 of the 7 had associated hyperbilirubinemia, and one subject died due to multiorgan system failure with lactic acidosis and evidence of hepatic steatosis.[7] This clinical trial emphasizes the need for high fidelity pre-clinical DILI risk prediction. Primary human hepatocyte (PHH) cell cultures currently used in these assays retain hepatocyte function but decline rapidly in metabolic function and vary greatly between cadaveric fresh and cryopreserved samples.[8] PHH availability and source patient diversity is also limited preventing their use in large-scale screening for DILI-risk.[9]

To meet this challenge, we explored the use of human liver organoids (HLOs) as a more physiologically organotypic system for recapitulating DILI *in vitro*, with added adaptations for high-throughput screening. HLOs consist primarily of hepatocyte-like cells while also containing non-parenchymal stellate-like and Kupffer-like cells derived from the same individual donor.[10] In this study, we utilized a previously-developed protocol for derivation of HLOs from induced pluripotent stem cells (iPSC) [11] allowing for a personalized approach in assessing DILI based on iPSC donor selection. We adapted HLOs both for high-throughput drug screening in 384-well plates and for enhanced physiological fidelity in a liver-chip system (Emulate Bio)[12] previously used to successfully predict species-specific DILI with PHHs.[13,14]

Due to the complexity and inconsistency of DILI based on culprit drug modality, we developed an integrated multi-omics platform including biomarker/analyte detection, high content imaging-enabled phenotyping, and single-cell RNA sequencing to deliver a comprehensive platform for dissection of DILI with inarigivir + tenofovir as a benchmark. In this study, we demonstrate the potential of dispersed HLOs for rapid 384-well based compound DILI-risk screening, and also the validation of a patient-derived liver-on-chip (PaDLOC) system for a more intricate and mechanistic assessment of DILI pathogenesis.

## EXPERIMENTAL PROCEDURES

### PaDLOC Culture and Compound Treatment

Dispersed HLO cells were transferred to the Chip S1™ based on the co-culture Liver-Chip Culture Protocol as described by Emulate Bio.[15] In brief, both channels were coated with an extracellular matrix consisting of collagen I (100 µg/mL) (Corning, 354249) and fibronectin (25 µg/mL) (ThermoScientific, 33010018) at 4 °C overnight followed by 1 hr at 37 °C. Dispersed HLOs were concentrated to a density of 4 × 10^6^ cells/mL. 50 μL of cell suspension was quickly pipetted into the bottom channel and the chip immediately flipped over to allow attachment to the membrane to allow even dispersion and cultured static (no media flow) at 37 °C for 8 hrs for attachment to the semi-permeable membrane. Next, 30 μL of cell suspension was seeded into the top channel and again left to attach for 8 hrs at 37 °C. Both channels were then washed with hepatocyte growth media containing hepatocyte growth factor, oncostatin M, dexamethasone de-gassed with a 0.45 μm Steriflip-HV Sterile Centrifuge Tube Top Filter Unit (MilliporeSigma, SE1M003M00).

Each seeded Chip S1 was then attached to a respective Pod™ Portable Module. Loaded Pods were placed into the Zoë™ Culture Module at 37 °C. All chips then underwent a regulate cycle followed by a constant flow rate of 30 µL/hr of the reservoir’s media for each of both channels modulated by an Orb™ Hub Module. Media outflow collected in respective reservoirs was obtained for ALT, AST, and albumin measurement. Fresh degassed hepatocyte growth media was added into the inlet reservoirs every 2 days. Cells were cultured with flow for 7 days before treatment.

After 7 days, residual hepatocyte growth media was aspirated and replaced with hepatocyte growth media containing either 0.1% DMSO, APAP (100 µM), FIAU (1 µM), tenofovir (500 nM), inarigivir soproxil (500 nM), or tenofovir and inarigivir soproxil (250 nM and 250 nM). The flow rate was maintained at a constant rate for an additional 7 days and outflow media was collected at days 1, 4, and 7 post-treatment.

### Dispersed 384-well HLO culture and drug delivery

Dispersed HLOs were seeded in collagen type 1-coated CellCarrier-384 Ultra Microplates (PerkinElmer, 6057308) at a seeding density of 8,000 cells/well in hepatocyte growth media. Cells were left to adhere and culture for 48 hrs before treatment with compounds. For screening, compounds were dispensed in 10-point dose-response from 2 nM to 500 μM using an HP D300e Digital Dispenser. For tenofovir-inarigivir synergy assessment, tenofovir, inarigivir sorpoxil, and in combination were dispensed in triplicate with 12-point dose-response curves in 1/3 dilutions starting with a high of 500 µM. Cells were then incubated for 120-hours before fixation and staining.

## RESULTS

### Use of Dispersed HLOs in 384-well Based High-Content Screening and Drug Clustering

Foregut spheroids from iPSC lines 72.3[16] (male fibroblast derived), 2E[17] (female fibroblast derived), and CC3[18] (female fibroblast derived) were generated through a 6-day differentiation and subsequently differentiated to HLOs over a 20-day period.[10,19] 3D confocal immunofluorescence imaging shows positivity for HNF4A to identify hepatocytes, α-SMA to identify stellate cells, and CD68 to identify Kupffer cells (Fig. S1A-B). Here, we dispensed 3,500 dispersed HLO cells/well in 384-well plates and observed slow proliferation across 7-days while they retained cell-type-specific markers (Fig. S1 A-C) on day 7 of culture. CellProfiler 4.2.0[20] cell segmenting determined 64.6-75.2% were positive for HNF4A, 18.7-32.4% for α-SMA, 0.12-0.19% for CD68 and 2.93-5.91% for neither (Fig. S1C). These cell ratios match previously determined cell distribution through single-cell RNA sequencing (scRNA-seq).[10]

With confirmation of retention of cell type specific markers and ratios in a 384-well format as in intact HLOs, a collection of 12 DILI-associated drugs were screened through HLOs developed through three different iPSC lines to characterize the drug-induced perturbation of single-cells in response to 12 hepatotoxic drugs from 384-well plate screening. These 12 compounds showed dose-responsive loss of cell viability with IC_50_ values in nanomolar to low micromolar range (Fig. 1A) while neither immortalized hepatocellular carcinoma line Huh7 and dispersed definitive endoderm obtained at an earlier development stage exhibited overt cytotoxicity (Fig. S2).

**Fig. 1.**
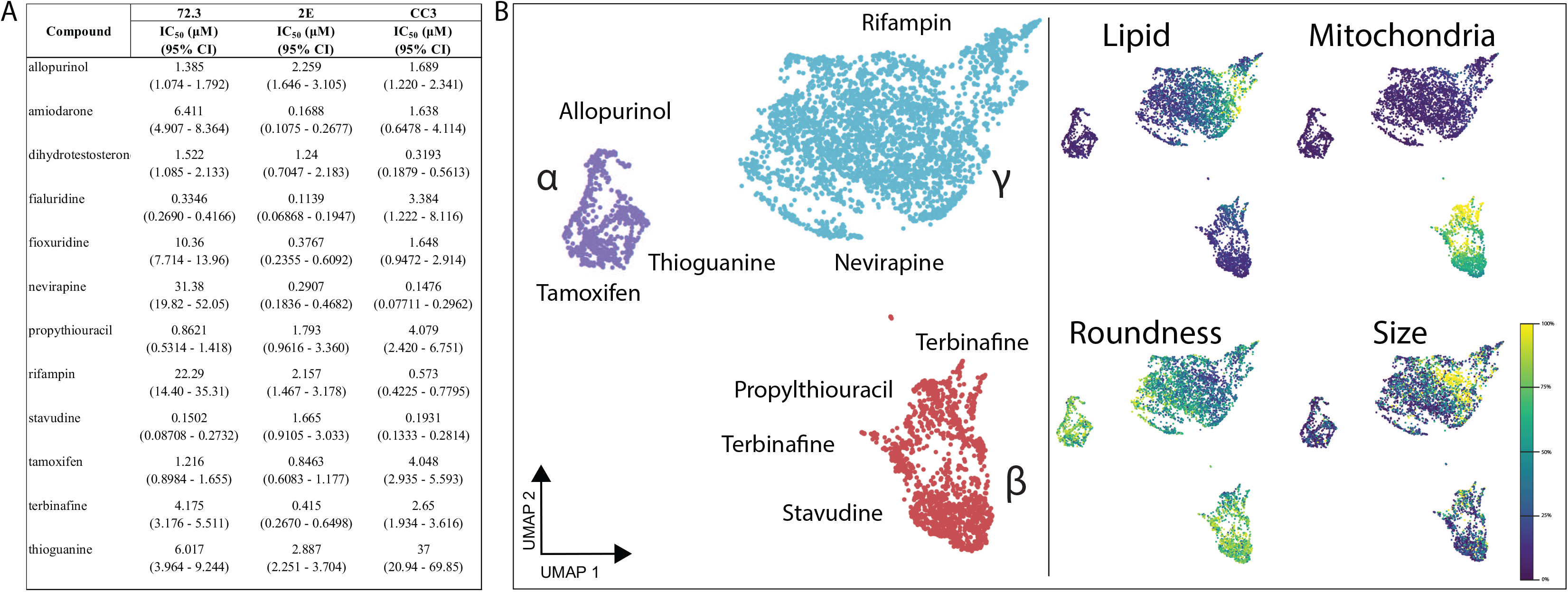
384-well adaptation of HLOs. HLOs grown from iPSC lines 72.3, 2E, and CC3 are dispersed into 384-well plates and treated with a 10-point dose response of 12 common DILI causing compounds. After 120 hrs incubation, cells are fixed and stained with Hoechst 33342, MitoView Green, HCS CellMaskOrange, and LipidTox DeepRed and imaged with an automated confocal microscope. (A) IC_50_ values of these compounds through cell viability counts are calculated (n=4 per concentration, per cell line). (B) CellProfiler was used to extract features at each compound’s respective IC_50_ values for 72.3-derived HLOs and embedded into UMAP. Plot points represent individual cells. Color intensity dictates the percentage of max measurement for each feature.

CellProfiler 4.2.0 was used to segment and extract 845 features per cell to generate a cell-by-feature matrix characterizing drug-induced perturbation of single-cells in response to the 12 hepatotoxic drugs. The dimensionality of the feature vector was reduced to 2-dimensions using the UMAP method[21] and hierarchical density-based clustering was performed with HDBScan[22] to characterize and cluster drug treatments by their resulting phenotypic perturbation. We observe three distinct clusters within this embedding (Fig. 1B, Fig. S3). Cluster α consists largely of allopurinol, tamoxifen, and thioguanine-treated cells. Cluster β largely contains cells treated with nucleotide/nucleoside analogs and consists mainly of cells treated with propylthiouracil, and to a lesser extent, stavudine, and thioguanine-treated cells. Lastly, Cluster γ contains a majority of cells treated with allopurinol and tamoxifen as in Cluster β, but with a major presence of nevirapine and rifampin which are thought to cause DILI through CYP450 modulation.[23] With other compounds, less pronounced clustering was observed (Fig. S3).

### Biochemical, phenotypic, and transcriptomic analysis of HLOs in an Organ-on-a-Chip System - iPSC Liver Chips

Patient-derived liver-on-chips (PaDLOC) were developed by dispersing HLOs into a single-cell suspension seeded in both upper and lower compartments of a dual-compartment microfluidic S1 chip (Emulate Bio) and cultured for an additional 7 days. The dual-compartment microfluidic S1 chip was previously used for long-term culture and maintenance of primary hepatocytes[24]. Primary hepatocytes on this system were shown to respond to DILI-causing compounds and recapitulate species-specific toxicity over the current preclinical standard models.[13]

While intact HLOs cultured on plates produce under 10 μg/mL albumin per day per 10^6^ cells with slight diminishment after 7 days of culture, PaDLOCs show an increased albumin production rate of 20-30 ug/mL per day per 10^6^ cells (Fig. 2A). This is comparable to albumin production by PHHs (Gibco, Lot HU8305) on this system and with previous findings while PHHs in plates quickly lost albumin production after few days in culture.[13] PaDLOCs also express CYP450s 1A1, 2D6, and 3A4 at 3-5 fold higher levels as compared to intact HLOs (Fig. S4). Increased CYP expression was also observed by metabolic turnover, as PaDLOCs metabolized acetaminophen (APAP), cyclophosphamide, and darunavir at increased rates compared to plate-cultured intact HLOs (Fig. 2B), albeit at slightly lower rates than PHHs.

**Fig. 2.**
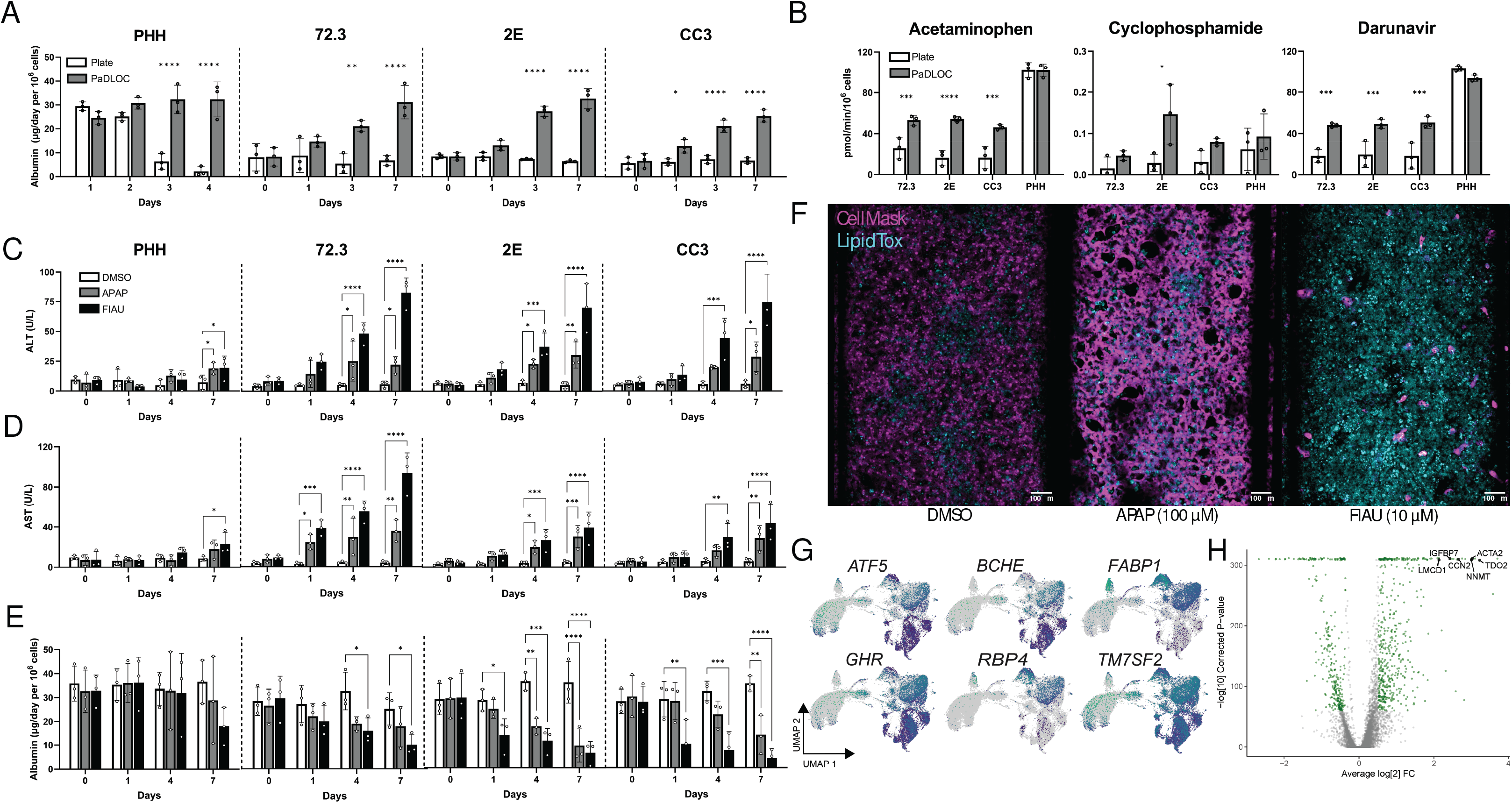
Development of a HLO-based Liver Chip. HLOs developed from iPSC lines 72.3, 2E, and CC3 are disrupted into single-cell suspension and cultured into patient-derived liver organoids on chip (PaDLOCs) and compared against intact organoids on 12-well plates. (A) Albumin released in PaDLOCs is identical to that of plate HLOs at day 0 but increases over 7 days (day 21-28 of differentiation). (B) PaDLOCs turnover CYP1A, 2B, and 3A family substrates acetaminophen, cyclophosphamide, and darunavir at increased rate compared to plate HLOs. (C) Cells are treated with DMSO control and known hepatotoxins APAP (100 μM) and FIAU (1 μM). PaDLOCs demonstrated both ALT (D) and AST release and (E) albumin production diminishment across 7 days. Bars and plot points represent mean ± SD (n=3 PaDLOC chips and n=3 plate HLO wells). Statistical significance was calculated using ANOVA with multiple comparison Dunnett’s test. *, **, ***, and **** denote P values of less than 0.05, 0.01, 0.001, and 0.0001 respectively. (F) Confocal images of PaDLOCs at day 7 of treatment stained with CellMask Orange (magenta) and LipidTOX Deep Red (cyan). Images shown are scaled to identical intensity ranges. (G) UMAP clustering of 72.3-derived PaDLOCs highlighting a selection of liver specific genes. Each point represents one cell. Gray values represent no detected expression. (H) Volcano plot comparing gene differential expression between 72.3-derived PaDLOC and HLOs with genes most upregulated in PaDLOC highlighted (>0 designates higher expression in PaDLOCs).

To confirm the presence of hepatocytes in PaDLOCs and compare the transcriptomic changes imparted by the chip system, we performed scRNAseq of iPSC 72.3 derived HLO cells using the Illumina NovaSeq platform. Single-cell analysis was performed for comparisons of culture conditions (Fig. 2G). Differential expression analysis between all cells of HLOs and PaDLOCs (Fig. 2H) showed an increase of liver proliferation biomarkers TGFBI (collagen binding) and CCN2 expression (cell adhesion) in PaDLOCs.[25] Increased expression of hepatocyte-marker TDO2, commonly correlated with increased CYPs 1A1 and 1A2[26] and ACTA2, a marker for activated stellate cells[27], were observed. Other liver-specific markers demonstrating increased expression in PaDLOCs include NNMT,[28] and IGFBP7.[29]

### iPSC liver chips for DILI risk prediction

Serum biomarkers for DILI include elevated ALT and AST[30] and diminished production of albumin.[31] These biomarkers correspond to hepatocellular injury. APAP and filauridine (FIAU) were chosen as compounds with known intrinsic hepatotoxicity with differing mechanisms of action. APAP is metabolized by CYP450-mediated oxidation to NAPQI which exerts hepatotoxicity via the formation of covalent liver protein adducts at cysteine residues as 3-(cystein-*S*-yl)-APAP.[32,33] In contrast, FIAU causes hepatotoxicity by stimulating ectopic lipid accumulation and as a mitochondrial toxin. FIAU infamously passed pre-clinical assays but still resulted in overt patient hepatotoxicity.[34,35] For all PaDLOC lines, treatment with 100 μM of APAP increased ALT from a basal level of less than 10 U/L at day 0 to peaks of around 20-30 U/L (Fig. 2C-D), while treatment with 10 μM FIAU drastically increased both ALT and AST to over 80 U/L. Additionally, albumin production, a guiding biomarker for the diagnosis of DILI severity,[31] was stable in DMSO-treated PaDLOCs while its production was diminished in both APAP and FIAU-treated PaDLOCs. These observations of ALT/AST release and reduced albumin production were not significant in PHH PaDLOCs (Fig. 2E).

The heterogeneity of DILI presents a challenge for DILI-risk prediction for novel therapeutics. For example, in patients, DILI from APAP and FIAU manifest differently due to differing mechanisms of action. APAP has been reported to cause hepatic necrosis[36] whereas FIAU causes diffuse microvesicular steatosis with retention of hepatic architecture.[37] PaDLOCs treated with APAP and FIAU at 100 µM and 10 µM, respectively, were stained for nuclei/cell regions and lipid droplets to test if PaDLOCs can capture this heterogeneity. Confocal images demonstrate APAP-treated PaDLOCs showed a patchy loss of cell mask and shriveling of cells with no increased lipid accumulation as compared to control (Fig. 2F). In contrast, FIAU treated PaDLOCs showed high lipid content and a reduction of CellMask staining.

### Modeling Hepatotoxicity of Tenofovir and Inarigivir Combinations

Cells in 384-well plates were treated with a 16-point dose range of tenofovir, inarigivir soproxil, and tenofovir - inarigivr in combination. After 120 hrs of treatment confocal images were taken for each treatment condition (n = 4 wells) stained to delineate nuclei/cell regions. Striking, while we observed negligible cytotoxicity for the monotherapies up to concentration of 100 μM, we observed 100% loss of cell viability in the tenofovir-inarigivir combination (Fig. S5A, IC_50_ = 56.9).

In PaDLOCs, tenofovir and inarigivir monotherapy at a concentration near their reported *C*_max_ (500 nM)[38,39] did not increase ALT or AST release after treatment and no morphological deviation from DMSO-treated control (Fig. 3). However, the combination of tenofovir and inarigivir increased ALT starting at day 4 to 15-25 U/L, and to 25-35 at day 7 and AST to 20-30 U/L at day 4 and 40-50 U/L at day 7 (Fig. 3A-B). Combination treatments also resulted in a decrease in albumin production while no effect was observed in the single-agent treatments across the 7 days (Fig. 3C). PaDLOCs from iPSC 72.3 did however demonstrate slight increase in ALT release only at day 7 of treatment.

**Fig. 3.**
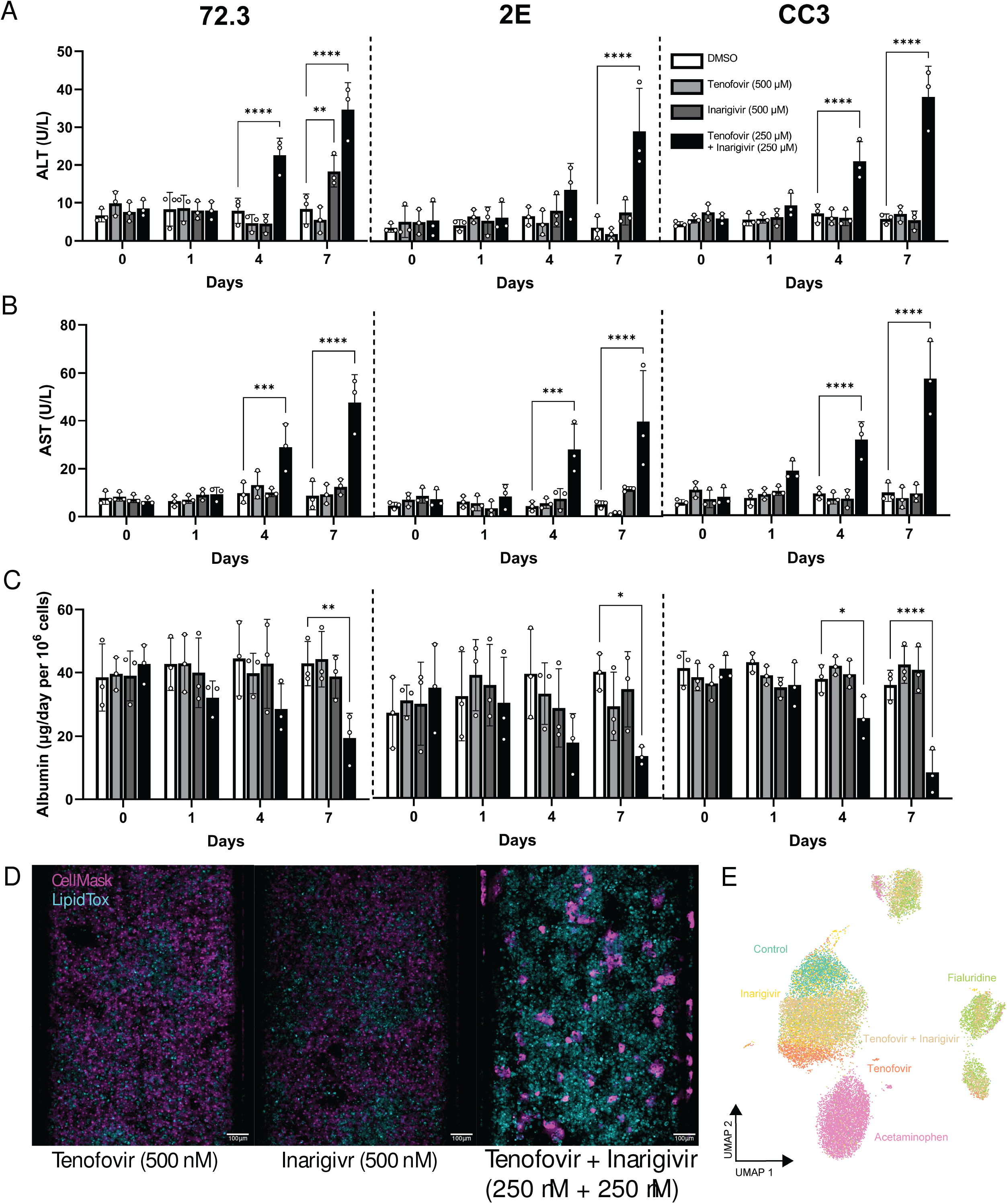
Assessment of known DILI-causing drug combination: tenofovi-inarigivir. (A) ALT, (B) AST, (C) and albumin released by 72,3, 2E, and CC3 PaDLOCs over 7 days of treatment with tenofovir (500 nM), inarigivir soproxil (500 nM), and tenofovir-xf inarigivir combination (250 + 250 nM) (n=3 chips per condition). Plot points represent mean ± SD. Statistical significance was calculated using ANOVA with multiple comparison Dunnett’s test. *, **, ***, and **** denote P values of less than 0.05, 0.01, 0.001, and 0.0001 respectively. (D) PaDLOCs treated with DMSO control, individual agents, combinations, APAP, and FIAU were stained with Hoechst 33342, CellMask Orange, and LipidTOX Deep Red. Images shown are scaled to identical intensity ranges. (E) CellProfiler extracted cell-level features were embedded into UMAP demonstrating morphological clustering. Plot points represent individual cells.

Visually, PaDLOCs treated with both combinations exhibited a similar phenotype to FIAU-treated controls with regional loss of CellMask staining and high lipid accumulation (Fig. 3D). These features were measured from confocal images and reduced via UMAP into a 2-dimensional projection (Fig. 3E). We note tenofovir-inarigivir-treated cells clustering with those FIAU-treated, while tenofovir and inarigivir single-agent treatments clustered with DMSO control. APAP treated cells exhibited a phenotype unlike either of the other groups and resulted in their unique cluster.

### Transcriptomic analysis of Tenofovir-Inarigivir, FIAU, APAP treated PaDLOCs

scRNA-seq was performed on drug-treated and control PaDLOCs on the Illumina NovaSeq platform. The concentration and duration of treatment were optimized using phenotypic endpoints to capture intermediate phenotypes rather than late-stage cell death. We evaluated DMSO-treated controls, fialuridine (10 μM), tenofovir (500 nM), and tenofovir-inarigivir (250 + 250 nM) combination. Single-cell data was subset to the hepatocyte population based on known hepatocyte markers as listed in Fig. 2G. Although tenofovir-inarigivir-induced hepatotoxicity has similar clinical features to that of fialuridine, our comparisons suggest a greater transcriptomic similarity between tenofovir monotherapy and fialuridine. First, UMAP re-embedding of the hepatocytes shows close clustering between fialuridine and tenofovir treatments (Fig. 4A). Volcano plots (Fig. 4C) show that both conditions, compared to control, result in overexpression of KCNQ10T1, upregulation of which was previously shown to diminish DILI effect[40] and suppressed expression of RPS10.

**Fig. 4.**
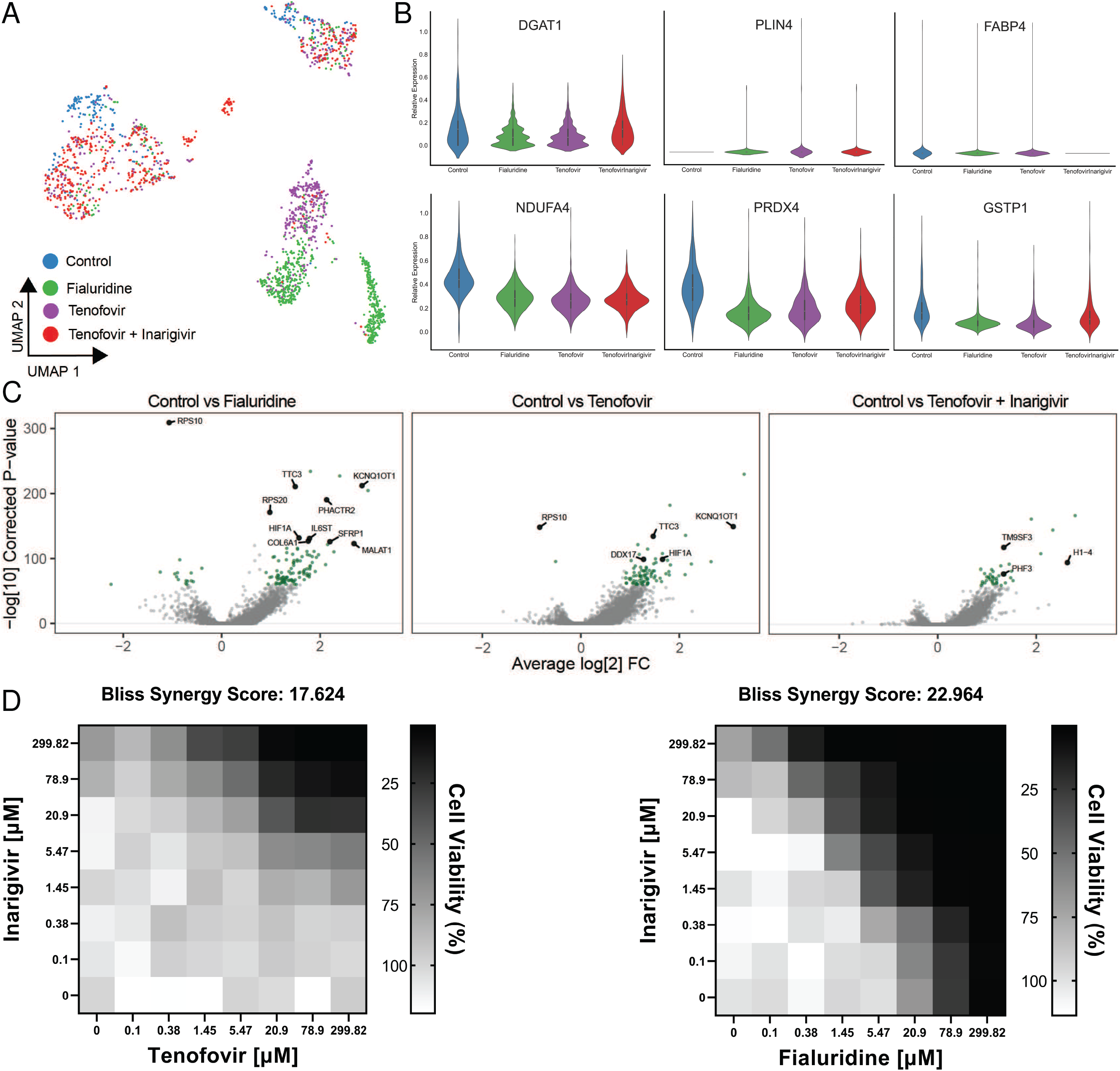
Single Cell Transcriptomics of treated PaDLOCs. (A) Hepatocytes across treatments are identified and subset through marker expression and embedded into a UMAP to visualize similarities between treatments. Plot points represent individual cells. (B) Relative expression of *DGAT1, PLIN4, FABP4, NDUFA4, PRDX4*, and *GSTP1* in vehicle control, fialuridine, tenofovir, and tenofovir-inarigivir treated PaDLOCs. (C) Volcano plots highlighting significant differential expression between control and drug treatments (>0 designates higher expression in treatment). (D) In 384-well cultures, 2-dimensional dose response assays show inarigivir and both tenofovir and fialuridine are synergistic with Bliss scores of 17.624 and 22.964, respectively, as calculated by SynergyFinder 2.0.

Hepatocyte-specific differential expression analysis shows correlative transcriptomic perturbation by fialuridine and tenofovir. Fatty acid, triglyceride, and lipid storage markers (Fig. 4B) were of particular interest due to observations through confocal microscopy (Fig. 3B). DGAT1, involved in triglyceride synthesis and storage,[41] is downregulated under both fialuridine and tenofovir treatment when expression under tenofovir-inarigivir combination treatment matches that of vehicle control. PLIN4, thought to aid in lipid droplet accumulation in the liver,[42] is not detected in the vehicle control but increased in all other conditions. FABP4 is expressed consistently in all but the combination treatment, conflicting with previous evidence that FABP4 is overexpressed in liver injury due to hepatocellular carcinoma.[43–45] However, other reports have shown that FABP4 knockdown results in greater adiposity in mice.[46] Across all treatments, we observe diminishment of NDUFA4 as compared to control.[47] We also observed decreased expression of PRDX4 in all treatments, and GSTP1 in fialuridine and tenofovir single treatments, indicators of oxidative stress.[48,49]

As we observed similar transcriptomic perturbations between fialuridine and tenofovir treatments, we assessed whether fialuridine-inarigivir combination treatments showed synergistic toxicity similar to tenofovir-inarigivir treatments. Interestingly, in 384-well dispersed HLO assays, both tenofovir-inarigivir and fialuridine/inarigivir combinatory treatment likely result in synergistic toxicity with calculated Bliss synergy scores of 17.624 and 22.964, respectively (Fig. 4D).

## DISCUSSION

HLOs both from our findings and other reports[50] show promise as a viable *in vitro* model for DILI risk prediction. They are amenable to both a high-throughput screening and adaptation to PaDLOCs to further enhance their organotypic function. Compared to PHHs, HLOs enable large-scale and high throughput DILI risk assessment due to their relative scalability and consistency. Their application in 384-well format serves as basis for an early-stage preclinical assessment of novel drugs. PaDLOCs exhibit physiological similarities to human liver including: 1) production of cell types from the same host genetics including hepatocyte-like cells, stellates and Kupffer cells, 2) albumin production, and 3) cytochrome P450 expression.

HLOs as a dispersed monolayer can minimize well to well variability and obtain clear single-cell resolution images. This enables high-content screening, Cell Painting,[51] and morphological cell profiling. As we demonstrated, these multivariate outputs cluster drugs by their phenotypic perturbation to infer similar mechanisms of action and compare to other compounds. Multivariate analysis also enables cumulative hepatotoxic scores unable to be defined by individual endpoints. For example, FIAU treatment at sub-cytotoxic concentrations results in multidimensional perturbation of cells including diminished mitochondrial mass and lipid accumulation (Fig. S5 B-C). Machine learning-based multivariate analysis can combine multiple features into a robust prediction score.

While high-throughput screening with dispersed HLOs in 384-well plates allows rapid expansion for large screening efforts across multiple iPSC lines, they suffer from lower CYP450 expression and lack of crucial hepatocyte function.[11] Although superior to hepatocellular carcinoma cell lines which often do not demonstrate hepatotoxicity, some observed IC_50_ values for HLOs loss of cell viability are higher than achievable *in vivo* C_max_. For example, the reported IC_50_ for nevirapine in 72.3 derived HLOs is over double that of reported C_max_ in patients.[52] PaDLOCs, likely due to more adequate drug metabolism and mixture of parenchymal and non-parenchymal cell types, seem to respond to many drugs at *in vivo* concentrations.

In our studies, fialuridine and tenofovir-inarigivir were administered at *C*_max_ concentrations and responded with clear hepatotoxicity across multiple cell lines. Hepatotoxicity of both therapies was not detected until clinical trials nor detected in the 384-well platform until higher concentrations. In APAP treatments, previous PHH studies on the Emulate system used 30-fold higher APAP concentrations to achieve a hepatotoxic effect[13], due to reliance of APAP turnover to NAPQI by CYP2E1.[53] Despite greater CYP expression and metabolic turnover, these effects are not observed at the same concentrations in PHHs suggesting also the necessity of the diverse cell types found in intact HLOs and PaDLOCs. Lastly, in patients, APAP and FIAU damaged liver histology present as hepatic necrosis[36] and diffuse microvesicular steatosis with retention of hepatic architecture,[37] respectively. Our confocal images show APAP-treated PaDLOCs with patchy loss of cell mass while FIAU-treatment results in over accumulation of lipids, seemingly mimicking their presentation in patient histology.

The integration of scRNA-seq, shown here as a proof-of-concept for liver chip systems, provides detailed predictive power for synergistic DILI. Herein, our unified multi-omics platform supported with transcriptomics data predicted FIAU-inarigivir synergy given known tenofovir-inarigivir synergy in a complex PaDLOC system, which was then confirmed in 2-dimensional dose response in the higher throughput platform. Although further optimizations are necessary, PaDLOCs show promise as a detailed model for DILI allowing multi-omic endpoints. As they are iPSC-based, and iPSCs can be reprogrammed from patient cells acquired non-invasively (e.g. PBMCs)[54] or even hESCs (Fig. S6), this platform can be expanded to encompass patient genetic diversity. An adequate biobank of PaDLOCs would be ideal to benchmark compounds before clinical trials and mitigate rare hepatotoxic events. Future studies will focus on developing a biobank of complex HLO co-cultures established from idiosyncratic DILI patients thereby concentrating genetics to a screenable number of patient lines as a predictive platform that can capture DILI risk with and enrichment of 10^6^ over the general population.

## Abbreviations

ALT: alanine aminotransferase
APAP: acetaminophen
DILI: drug-induced liver injury
FIAU: fialuridine
HLA: human leukocyte antigen
HLO: human liver organoid
iPSC: induced-pluripotent stem cells
PaDLOC: patient-derived liver-on-chip
PHH: primary human hepatocytes
scRNA-seq: Single cell RNA sequencing
UMAP: Uniform Manifold Approximation and Projection

## Acknowledgements

The authors would like to thank Emulate Inc. for assistance with experimental design. We thank Kevin Jan at Yokogawa for microscopy support. We thank Teresa O’Meara for writing assistance and proofreading, and Carmen Mirabelli, Chung Owyang, and Bishr Omary for thoughtful advice. We thank Andrew Tidball, Wei Niu, and Xiaotian (Tracy) Qiao for gifting the iPSC lines and providing cell culture support. Lastly, we thank Yihao Zhuang for assistance with acquiring LC/MS data.

## Data Availability

All relevant data is provided within this paper and its Supplementary Information. PaDLOC scRNA-seq data have been deposited in the Gene Expression Omnibus (GEO) under accession number GSE188541.

## Supporting Information

Supplementary Materials and Methods

Fig. S1

Fig. S2

Fig. S3

Fig. S4

Fig. S5

Fig. S6

Author names in bold designate shared co-first authorship.

## Graphical Abstract

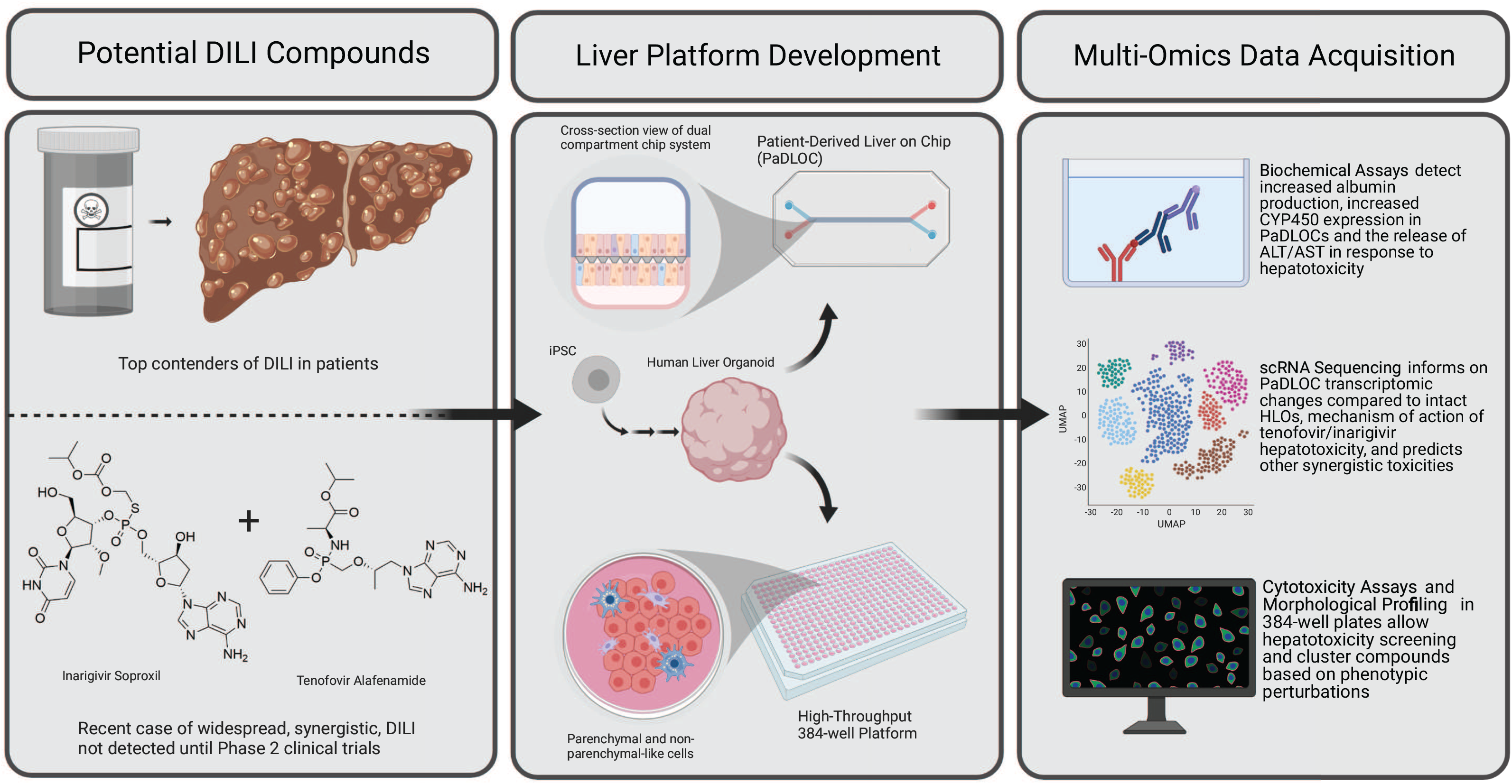

## Highlights

- Liver organoids retain functionality in a high-throughput platform
- Phenotypic clustering identifies similarities in drug mechanisms of hepatic injury
- Organoids on chip increase albumin and CYP expression; respond to hepatotoxic drugs
- Liver chips model hepatotoxicity of tenofovir-inarigivir combination therapy
- Transcriptomics with morphological profiling predict other synergistic toxicities

## Notes

**Conflicts of interest:** RJF has research support from Gilead and Abbvie. Other authors have no conflicts.

### Competing Interest Statement

The authors have declared no competing interest.

### Summary of Updates

We have incorporated a total of 3 independ iPSC-derived liver organoid lines to enhance the rigor and reproducibility of the experimental system. The additional liver organoid lines recapitulated the drug-induced liver injury caused by tenofovir-inarigivir and show the utility of this approach toward DILI risk prediction. We also have performed additional experiments including metabolomic quantitation of drug metabolism as a measure of CYP450 activity and have improved the immunofluorescence imaging for cell type markers by selection and optimization of new antibodies.

