## Supplemental for "A Human Liver Organoid Screening Platform for DILI Risk Prediction"

### Supplementary Materials and Methods

*Human liver organoid culture and dispersion*

Human iPSC line 72.3 was obtained from Cincinnati Children's Hospital Medical Center[1] and iPSC lines 2E[2] and CC3[3] were gifted by the University of Michigan Human Stem Cell and Gene Editing Core. iPSCs were differentiated into HLOs based on a previously described protocol.[4,5] In brief, iPSCs were grown to 90% confluency on growth factor reduced Matrigel (Corning, 354230) coated 6-well plates (ThermoScientific, 140675). Cells were then treated with Activin A (R&D Biosystems, 338-AC) for 3 days and FGF4 (purified in house[6]) for 3 additional days to form definitive endoderm spheroids. Spheroids were embedded in 75 µL Matrigel (Corning, 354234) droplets in 24 well plates (ThermoScientific, 1142475) and treated with retinoic acid for 4 days followed by hepatocyte growth media (Hepatocyte Culture Medium BulletKit (Lonza, CC-3198) supplemented with 10 ng/mL hepatocyte growth factor (PeproTech, 100-39), 20 ng/mL oncostatin M (R&D Systems, 295OM050), and 0.1 µM dexamethasone (Millipore Sigma, D4902)) for 12 days.

HLOs were then taken out of Matrigel by treatment with dispase (0.2 mg/mL) for 10 minutes at 37 $^{o}$C followed by washing with DMEM/F12 (ThermoScientific, 11320033) and centrifugation at 300 x g for 3 minutes to pellet. To achieve a single cell suspension, cells were treated with trypsin (0.25%) (Invitrogen, 25200056) and incubated at 37 $^{o}$C for 10 minutes, mechanically dissociated by pipetting, and incubated until dissociation for up to an additional 10 minutes. Trypsin was quenched with 100% FBS (Corning, 35-010-CV) and washed 3 times with DMEM/F12 followed by resuspension in HCM.

*Plate and Organoid Fixation and Staining*

Plates were fixed with 4% paraformaldehyde for 15 minutes followed by permeabilization by 0.1% Triton X-100 (MP Biomedicals, 194854) for 15 minutes (50 µL per well). Cells were then blocked by a buffer containing 5% BSA (MilliporeSigma, A9647) and 0.01% Tween (FisherScientific, BP337-100) in PBS for 1 hour (50 µL per well). At this point, the protocol deviated based on necessary stains.

For 384-well hepatotoxicity assays, plates were stained with Hoechst 33342, MitoView Green (Biotium, 70054), HCS CellMask Orange (ThermoFisher Scientific, H32713), and HCS LipidTox Deep Red (ThermoFisher Scientific, H34477). PaDLOCs were stained with Hoechst 33342, HCS CellMask Orange, and HCS LipidTox Deep Red. Stains were diluted based on manufacturer's recommendations in blocking buffer and applied to cells (25 µL per well), washed 3 times with PBS (50 µL per well).

For 384-well biomarker immunofluorescence assays, cells were stained with Hoechst 33342 (ThermoFisher Scientific, H3570), HNF4A (ThermoFisher Scientific, MA5-14891), α-SMA (Abcam, ab21027), and CD68 (Abcam, ab53444). Antibodies were diluted at 1:500 in blocking buffer and added to each well and incubated overnight at 4 $^{o}$C (25 µL per well). Antibody solution was then washed 3 times with PBS (50 µL per well) and stained with a secondary solution. For immunofluorescence containing rat, rabbit goat, primary antibodies, the secondary solution contained anti-rat Alexa Fluor 488, anti-rabbit Alexa Fluor 555, and anti-mouse Alexa Fluor 647 Highly Cross-Adsorbed Secondary Antibodies (ThermoFisher Scientific, A21208, A31572, A31571). Plates were incubated for 1 hour at room temperature followed by another three PBS washes before imaging.

*Image Acquisition*

All images were acquired with a Yokogawa CQ1 Benchtop High-Content Screening System. 384-well plates were imaged with 1μm Z-spacing and 15 μm depth at 20X (Olympus UCPLFLN20X). PaDLOCs were imaged with 1 μm sections through 100 μm depth also at 20X.

*Cell Morphological Profiling and Data Analysis*

Individual nuclei were first delineated in the Hoechst 33342 stained channel using Cellpose 2.0.[8] Multi-channel fluorescence images were then analyzed with CellProfiler 4.2.0. Whole cell was delineated using nuclei as seed objects and dilation to the extent of the cell boundary which enabled measurements of fluorescent intensity, intensity distribution, texture, size, and shape from the respective regions in each fluorescent channel.

The resulting data set included hundreds of measurements on a per-cell level with necessary cell-level metadata. Cell viability across compound dose-range was obtained based on the number of identified cells per condition with DMSO vehicle-treated control as the 100% viability reference. Uniform Manifold Approximation and Projection (UMAP) embedding was done with the Python umap-learn package. Measurements for each feature were centered at zero and scaled with Z-score=1. Features with low variance were omitted.[9] Bliss synergy scores were calculated using SynergyFinder 2.0.[10]

*Cell Type Confirmation by Marker Positivity*

Based on visual inspection, cells positive for respective markers were selected to obtain an estimated intensity value for positivity in relation to cell compartment (nuclear for HNF4A, cytoplasmic for α-SMA and CD68) in CellProfiler Analyst 3.0[11] and set as thresholds for cell type identification. Cells with no expression of these three markers were classified as “other”.

*Single-Cell Transcriptomics*

HLOs were dispersed as described previously and cells were removed from control and treated PaDLOCs with trypsin (0.25%) (Invitrogen, 25200056) followed by FBS inactivation and PBS washing. Cells were kept on ice, confirmed to have >85% cell viability, and sent to the University of Michigan Advanced Genomics Core to prepare single cell libraries on the 10x Chromium with a cell capture target of 5,000 cells. Sequencing was performed by the core on a NovaSeq 6000 with a target of 100,000 reads per cell.

Each sample generated between 440 and 860 million barcoded reads corresponding. Transcripts were mapped to the GRCh38 2020-A (GENCODE v32/Ensembl 98) (July 7th, 2020) reference transcriptome[12] using 10x Genomics Cell Ranger 5.0.1,[13] where between 45% and 58% of reads per sample confidently map to the transcriptome, yielding between 500 and 4,700 median genes per cell. As a quality filter, genes were excluded if they were only detected in 5 or fewer cells, and cells were excluded if over 30% of reads were mitochondrial or if they had fewer than 10,000 total reads. Given that hepatocytes may be binucleated,[14] we did not remove doublets.

To estimate differential expression, we first correct for over-dispersion using sctransform v2 [15] with Seurat v4[16] which fits a robust negative binomial model for the per-gene variance by the expression mean. We then used DESeq2 to compute the average log fold-change and adjusted p-value for each gene between cells under different conditions.[17]

*UMAP Embedding*

To visualize and cluster cells with distinct phenotypes, we used UMAP non-linear dimensionality reduction[18]. For scRNA-seq data, UMAP embeddings were done in monocle3.[19] Expression values were normalized by dividing by per-cell size factors, adding a pseudo-count of 1, and taking the natural log; reducing the dimensionality with principal component analysis to 100 dimensions; and then applying the UMAP algorithm implemented in uwot[20] using the monocle3 default arguments: similarity="cosine", min_dist=0.1, n_neighbors=15. For CellProfiler morphological data, cell features were normalized (mean=0, stdev=1) and cells were filtered down to n-500 cells for each treatment compound following embedding in umap-learn.[18]

*Human Serum Albumin, ALT, and AST Measurements*

Media from PaDLOCs outflow or culture plates were obtained. HSA from chip media outflow was measured using an albumin human ELISA Kit (ThermoFisher Scientific, EHALB). 10 uL of media was diluted 100-fold in PBS before incubation on ELISA plate. A standard curve was made in accordance with kit guidelines and used to determine albumin concentration in all samples. Each sample was assayed in triplicate.

For ALT and AST measurement, 30 uL of media and PBS blanks were dispensed into respective wells in a 96-well assay plate. 300 uL of room temperature ALT/GPT Reagent (Thermo Scientific, TR71121) or AST/GOT Reagent (Thermo Scientific, TR70121) was then dispensed into all wells in the plate and incubated at 37 $^{o}$C for 30 seconds before recording absorbance at 340 nm for 3 minutes. Activity of ALT or AST was determined using the following equation:

$$\text{Abs/min} \times \text{Factor}$$

Where Factor is pre-determined for this assay and is 1746, from the manufacturer's manual. Average activity from blanks was subtracted from all other samples. All plates were read with a BioTek Synergy H1 Microplate Reader.

*CYP450 Expression Quantification*

RNA was purified with the Direct-zol RNA Miniprep (Zymo Research, R2052) and expression was measured using CFX96 Touch Deep Well Real-Time PCR System (BioRaD) and iTaq Universal Probes One-Step Kit (BioRad, 1725141). Primers used were CYP1A1 (ThermoFisher Scientific, Hs01054796_g1), CYP2D6 (Hs04931916_gH), CYP3A4 (Hs00604506_m1), and housekeeping gene GAPDH (ThermoFisher Scientific, Hs02786624_g1). Fold change was calculated using the ΔΔCt method over PaDLOC.

*CYP450 Metabolic Turnover*

Acetaminophen, cyclophosphamide, and darunavir were chosen as substrates for CYP 1, 2, and 3 families respectively. For this purpose, PaDLOCs at day 7 were taken out of the Zoe Culture Module. Media in PaDLOCs and plate cultures were replaced with 500 µL media containing a cocktail of the substrates each at 10 µM. 100 µL of media was taken after 1 hour and 2 hours of incubation. Reactions were immediately stopped using 100 µL of cold methanol and centrifuged for 5-minutes at 3,000 RPM and supernatant was collected.

Samples and substrate standards (0.11, 0.33, and 1 uM in media:methanol 50:50) were measured using an Agilent qTOF 6545 LC/MS system with the following parameters: Phenomenex Kinetex 1.7µm Phenyl-Hexyl

100 Å (50 x 2.1 mm) column; 2 µL injection volume; LC gradient, solvent A, 5% acetonitrile and 0.1% formic acid; solvent B, 100% acetonitrile and 0.1% formic acid; 0 min, 0% B; 1 min, 0%B; 7 min, 80% B; flow rate 0.4mL/min; MS, positive ion mode. Quantification was done using Masshunter Workstation by peak area integration from extracted ion chromatograms (EIC) for target masses of 152.0706, 261.0321, and 548.2425 for acetaminophen, cyclophosphamide and darunavir, respectively. Linear regressions were fit in KNIME for each standard vs. peak area and used to determine concentration of all samples.

*Dose-Response Curve and Statistical Analysis*

Dose-response curves were fitted and $IC_{50}$ values were determined in GraphPad Prism 9 using the nonlinear regression [inhibitor] vs response method. Statistical significance was calculated using ANOVA with multiple comparison Dunnett's test. *, **, ***, and **** denote P values of less than 0.05, 0.01, 0.001, and 0.0001 respectively.

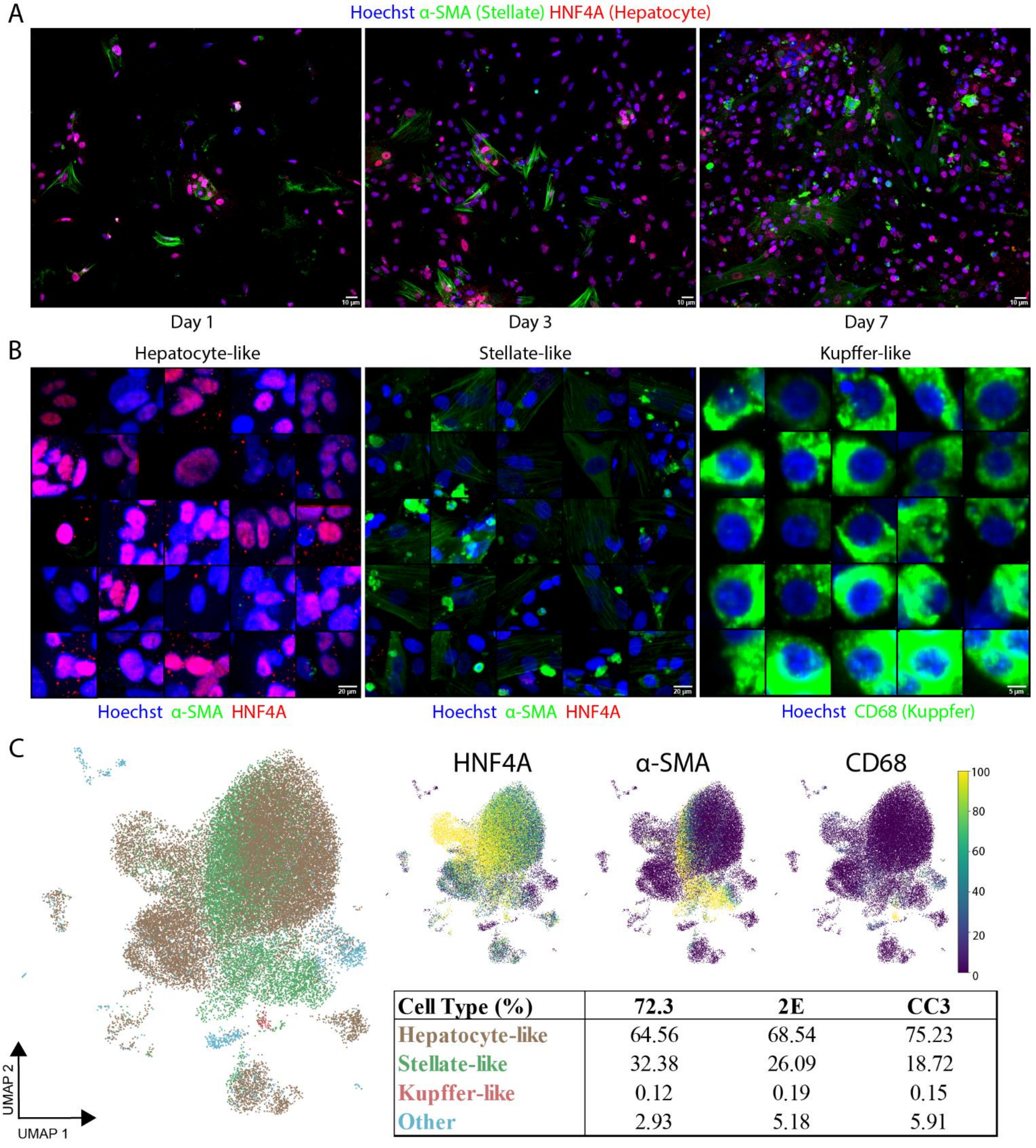


| Cell Type (%) | 72.3 | 2E | CC3 |
|---|---|---|---|
| Hepatocyte-like | 64.56 | 68.54 | 75.23 |
| Stellate-like | 32.38 | 26.09 | 18.72 |
| Kupffer-like | 0.12 | 0.19 | 0.15 |
| Other | 2.93 | 5.18 | 5.91 |

**Fig S1.** (A) Confocal images of 384-well monolayer cultures of dispersed HLOs across 7 days of culture showing retention of cell type specific markers HNF4A (hepatocytes) and α-SMA (stellates). (B) Collage of a subset of identified hepatocyte-like, stellate-like, and kupffer-like cells assembled in CellProfiler Analyst 3.0. (C) UMAP embedding of cell morphological features of 384-well monolayer cultures with the previous stain set. Percentage of hepatocytes, stellates, and Kupffer cells were estimated by marker positivity. Intensity scales of markers were determined from a range of empty background to highest cell intensities.

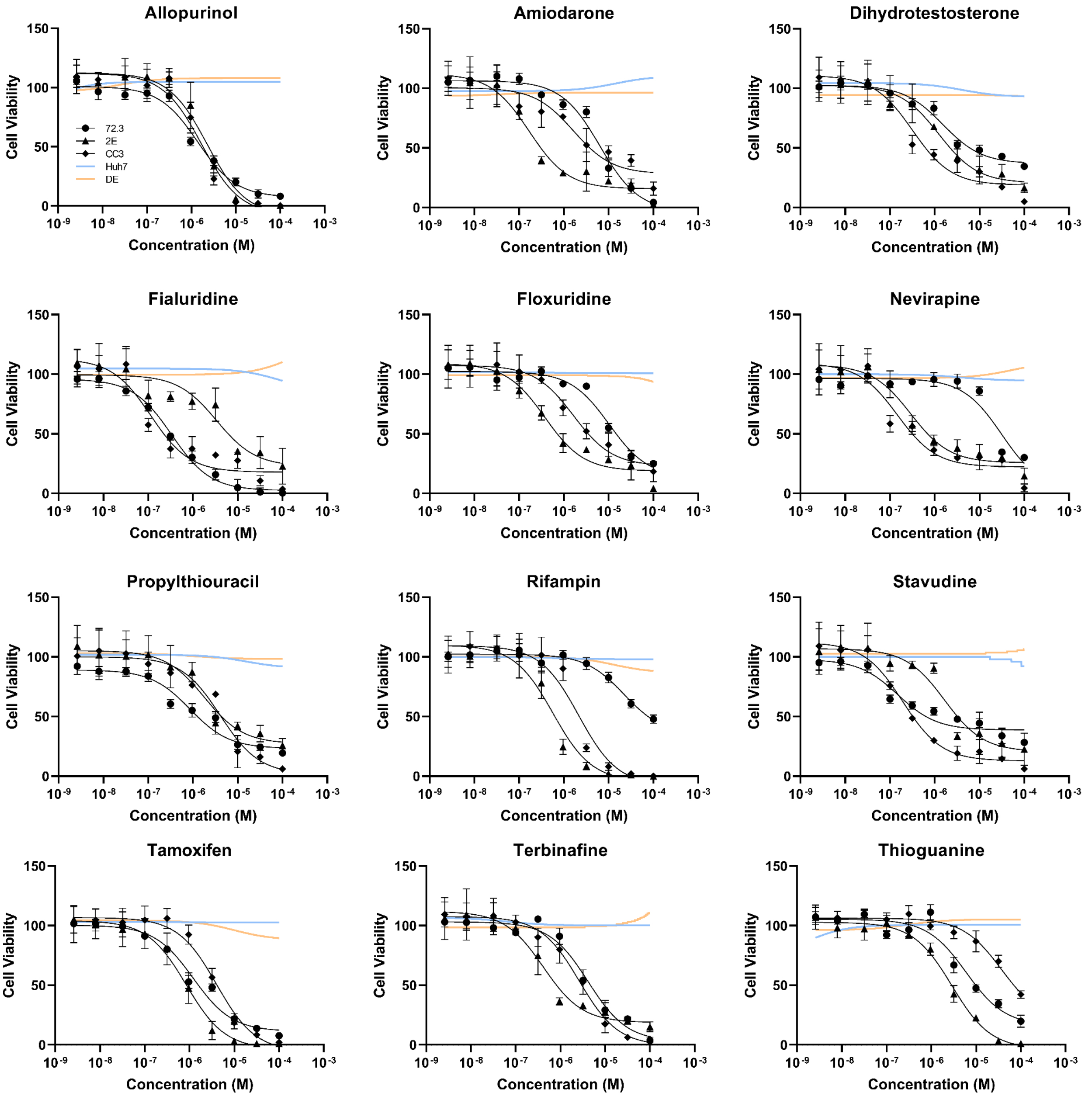


**Fig S2.** Cell viability dose response curves for 12 compounds commonly implicated in DILI against HLOs grown in three independent iPSC lines dispersed into 384-well plates and used to calculate $IC_{50}$ values shown in Figure 1. Immortalized cell lines and definitive endoderm from earlier in the HLO differentiation process are included as controls.

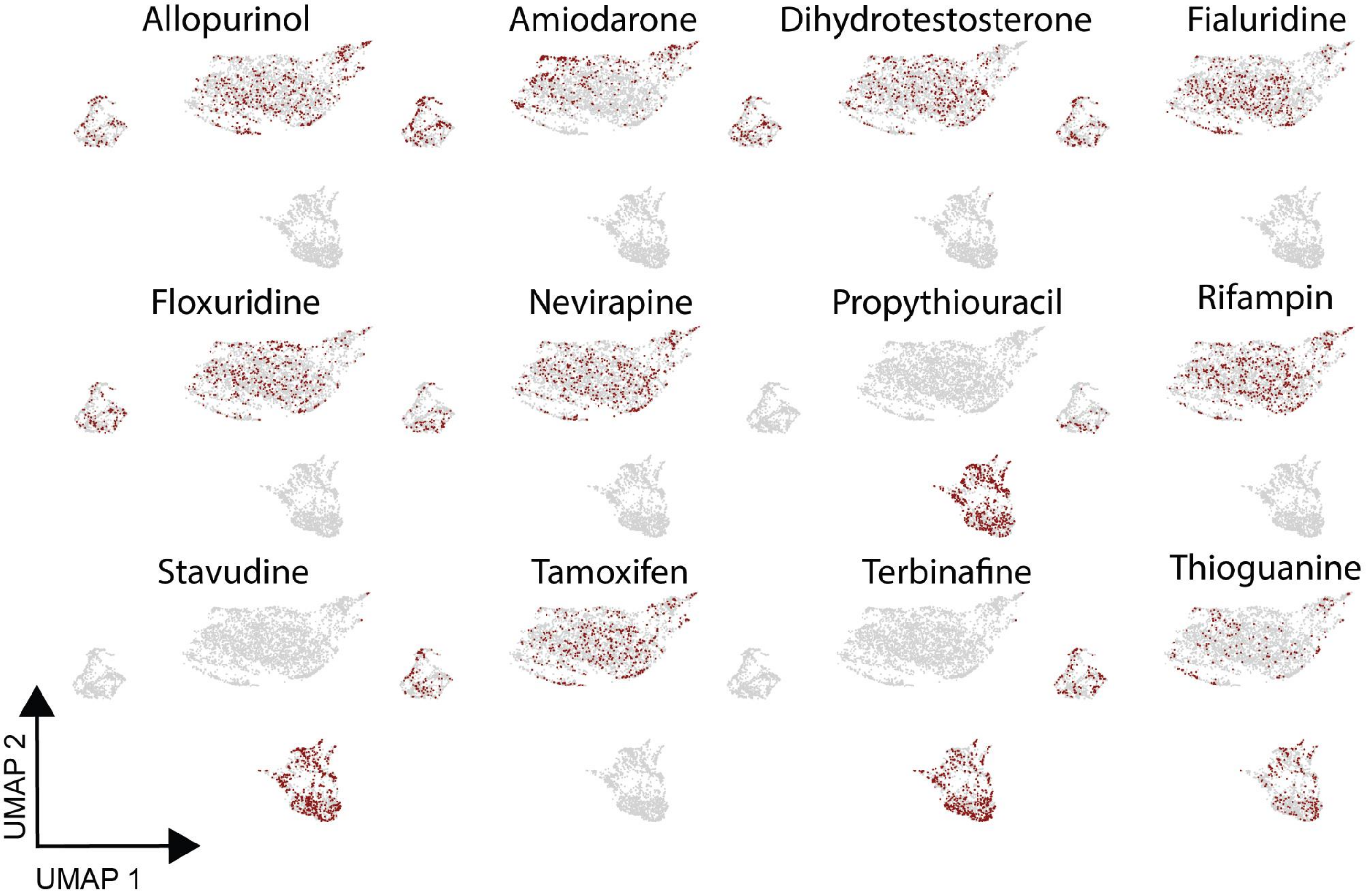


**Fig S3.** HLOs were dispersed into 384-well plates and treated with 10-point dose response for 12 commonly identified DILI compounds. Plates were then fixed and stained with Hoechst 33342, MitoView Green, HCS CellMask Orange, and HCS LipidTox Deep Red. CellProfiler 4.2.0 was used to extract morphological features of cells at their respective $IC_{50}$ values and embedded into UMAPs with respective compounds highlighted in red.

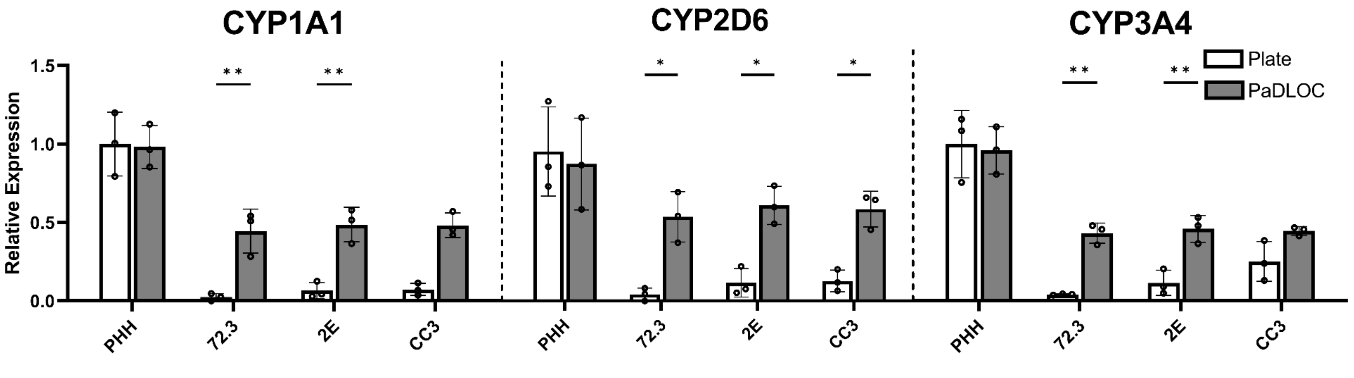

**Fig S4.** CYP 1A1, 2D6, and 3A4 expression of PHHs and HLOs grown from iPSC lines 72.3, 2E and CC3 on plate and after 7 days culture on PaDLOC. Statistical significance was calculated using ANOVA with multiple comparison Dunnett's test. * and ** denote P values of less than 0.05 and 0.01 respectively.

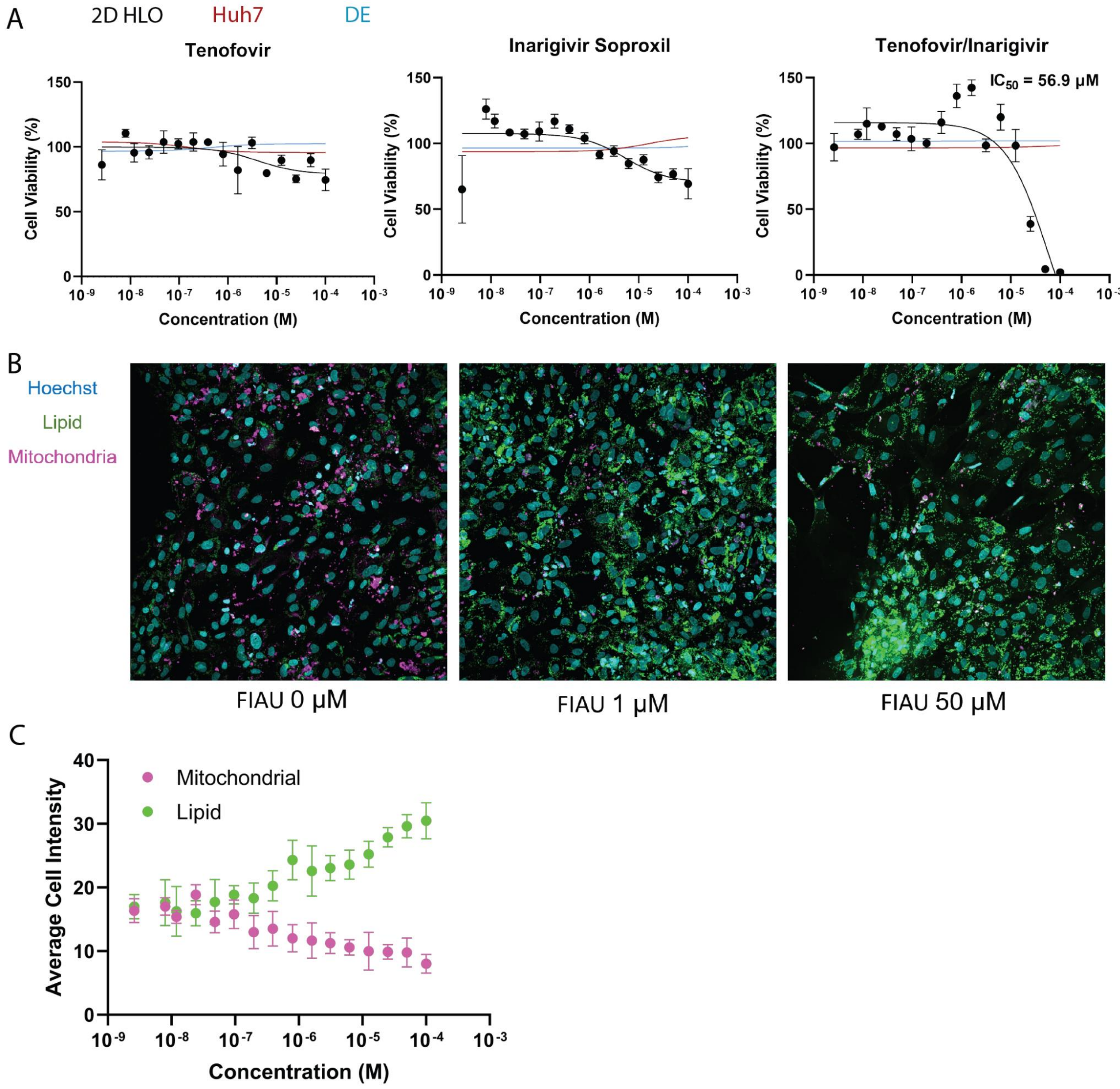


**Fig S5.** (A) Cell viability of 2D 384-well monolayer cultures of dispersed HLOs treated in 16-point dose-response with tenofovir, inarigivir soproxil, or in combination (n=4 per concentration) and measured $IC_{50}$. (B)

Confocal microscopy of FIAU treated 2D monolayers stained for nuclei, lipids, and mitochondria and (C) the per-cell measurement of these features across a dose-range of FIAU. Plot points represent mean ± SD (n=4 per concentration).

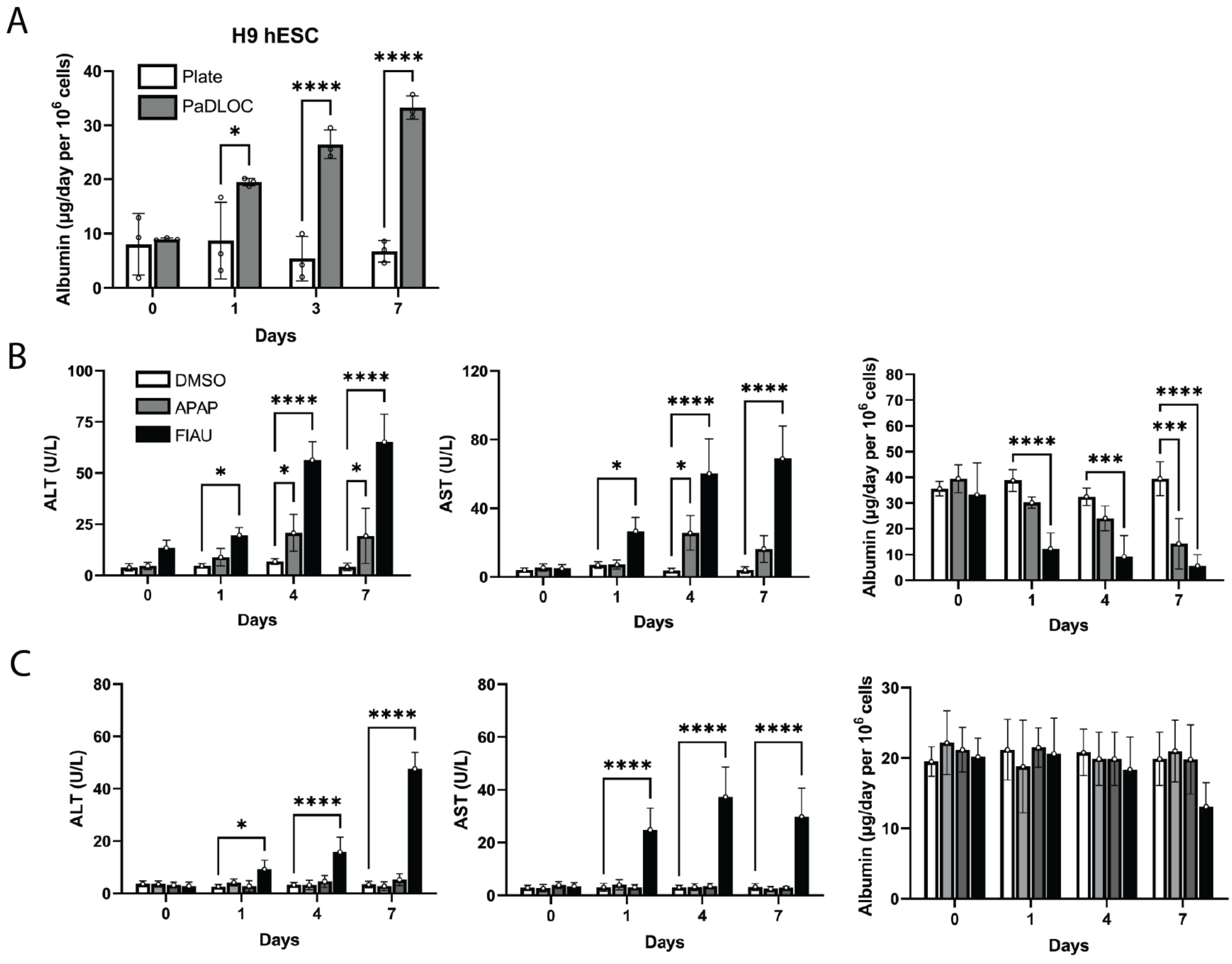


**Fig S6.** H9 human embryonic stem cell derived PaDLOCs show (A) increased albumin production as compared to intact HLOs and (B) consistent ALT, AST, and albumin response to known DILI compounds. (C) H9 PaDLOCs also respond to inarigivir/tenofovir induced hepatotoxicity without apparent hepatotoxicity in response to the individual compounds

[2] Dang LT, Vaid S, Lin G, Swaminathan P, Safran J, Loughman A, et al. STRADA-mutant human cortical

organoids model megalencephaly and exhibit delayed neuronal differentiation. Dev Neurobiol 2021;81:696–709.
[3] Tidball AM, Neely MD, Chamberlin R, Aboud AA, Kumar KK, Han B, et al. Genomic Instability Associated with p53 Knockdown in the Generation of Huntington's Disease Human Induced Pluripotent Stem Cells. PLoS One 2016;11:e0150372.
[4] Thompson WL, Takebe T. Generation of multi-cellular human liver organoids from pluripotent stem cells. Methods Cell Biol 2020;159:47–68.
[5] Ouchi R, Togo S, Kimura M, Shinozawa T, Koido M, Koike H, et al. Modeling Steatohepatitis in Humans with Pluripotent Stem Cell-Derived Organoids. Cell Metab 2019;30:374–84.e6.
[6] Sugawara S, Ito T, Sato S, Sato Y, Kasuga K, Kojima I, et al. Production of an aminoterminally truncated, stable type of bioactive mouse fibroblast growth factor 4 in Escherichia coli. J Biosci Bioeng 2014;117:525–30.
[7] Dekkers JF, Alieva M, Wellens LM, Ariese HCR, Jamieson PR, Vonk AM, et al. High-resolution 3D imaging of fixed and cleared organoids. Nat Protoc 2019;14:1756–71.
[8] Stringer C, Pachitariu M. Cellpose 2.0: how to train your own model. bioRxiv 2022:2022.04.01.486764. https://doi.org/10.1101/2022.04.01.486764.
[9] Becht E, McInnes L, Healy J, Dutertre C-A, Kwok IWH, Ng LG, et al. Dimensionality reduction for visualizing single-cell data using UMAP. Nat Biotechnol 2018. https://doi.org/10.1038/nbt.4314.
[10] Ianevski A, Giri AK, Aittokallio T. SynergyFinder 2.0: visual analytics of multi-drug combination synergies. Nucleic Acids Res 2020;48:W488–93.
[11] Stirling DR, Carpenter AE, Cimini BA. CellProfiler Analyst 3.0: Accessible data exploration and machine learning for image analysis. Bioinformatics 2021. https://doi.org/10.1093/bioinformatics/btab634.
[12] Frankish A, Diekhans M, Ferreira A-M, Johnson R, Jungreis I, Loveland J, et al. GENCODE reference annotation for the human and mouse genomes. Nucleic Acids Res 2019;47:D766–73.
[13] Zheng GXY, Terry JM, Belgrader P, Ryvkin P, Bent ZW, Wilson R, et al. Massively parallel digital transcriptional profiling of single cells. Nat Commun 2017;8:14049.
[14] Duncan AW, Taylor MH, Hickey RD, Hanlon Newell AE, Lenzi ML, Olson SB, et al. The ploidy conveyor of mature hepatocytes as a source of genetic variation. Nature 2010;467:707–10.
[15] Choudhary S, Satija R. Comparison and evaluation of statistical error models for scRNA-seq. bioRxiv 2021.
[16] Hao Y, Hao S, Andersen-Nissen E, Mauck WM 3rd, Zheng S, Butler A, et al. Integrated analysis of multimodal single-cell data. Cell 2021;184:3573–87.e29.
[17] Love MI, Huber W, Anders S. Moderated estimation of fold change and dispersion for RNA-seq data with DESeq2. Genome Biol 2014;15:550.
[18] McInnes L, Healy J, Melville J. UMAP: Uniform Manifold Approximation and Projection for Dimension Reduction. arXiv [statML] 2018.
[19] Cao J, Spielmann M, Qiu X, Huang X, Ibrahim DM, Hill AJ, et al. The single-cell transcriptional landscape of mammalian organogenesis. Nature 2019;566:496–502.
[20] Melville J, Lun A, Djekidel MN. uwot: The uniform manifold approximation and projection (UMAP) method for dimensionality reduction. R Package Version 2020;15.
